# Cryo-tides: synthetic peptides for cryo-EM grid preparation of multi-subunit protein complexes sensitive to the air-water interface

**DOI:** 10.64898/2026.09.30.755610

**Authors:** Rose Young, Amy J. S. McGeoch, Matthew Jessop, Basil J. Greber, Claudio Alfieri

**Affiliations:** Division of Structural Biology, Chester Beatty Laboratories, The Institute of Cancer Research, London, UK; The Francis Crick Institute 1, Midland Road, London UK

## Abstract

Cryogenic electron microscopy (cryo-EM) has enabled rapid progress in structural biology throughout the last decade. However, cryo-EM sample preparation still represents a major bottleneck in high-resolution structure determination. Many proteins and macromolecular complexes interact with the air-water interface during sample preparation, which can cause protein denaturation, complex disassembly, and preferred particle orientation. Development of methods to alleviate these issues is an active field of investigation. However, a universally applicable solution to this problem has not been found so far. Here, we describe the simple and cost-effective usage of synthetic peptides with defined amino acid sequence to protect cryo-EM samples from complex disassembly and reduce preferred orientation. We show benefits during cryo-EM grid preparation of a large E3 ubiquitin ligase termed the anaphase-promoting complex (APC/C), and the small CDK11:cyclin L:SAP30BP complex.

## Introduction

In the past decade, cryogenic electron microscopy (cryo-EM) has rapidly matured into a major method for the determination of high-resolution structures of biological macromolecules, routinely achieving resolutions of 4 Å or better (reviewed in^1–3^) and in favourable cases even exceeding 2 Å resolution (e.g. ^4–10^). Cryo-EM specimen preparation commonly involves the application of a small droplet (often 3-4 µL) of sample solution containing a biological specimen to a support grid. The liquid film is then thinned by blotting (often to below 50 nm thickness) and rapidly frozen by plunge-freezing in liquid ethane or a similar cryogen^11^. As a result of the rapid cooling process, the sample has no time to form ice crystals and forms electron-transparent vitreous ice^12–15^. This vitrification step is the key to the success of cryo-EM because it preserves the molecular complexes contained in the sample in a frozen-hydrated, near native state while allowing the specimen to endure the high vacuum inside an electron microscope. However, the need for formation of a thin liquid film just before freezing creates a large surface-to-volume ratio, a condition that can lead to damage of the macromolecular complexes in the sample^16^. Therefore, the sample preparation step remains as a main bottleneck that hinders rapid and reproducible structure determination of biological macromolecules by cryo-EM ^17,18^.

The key issue underlying these challenges is that many macromolecules interact with the air-water interface (AWI) during the grid freezing process^16,18^. It has been estimated that molecular complexes collide with the AWI or the grid support thousands of times before the grid is frozen^16^. Consequently, a systematic analysis found that particles are commonly adsorbed to the AWI in cryo-EM specimens^19^. In addition to damaging the sample, interactions with the hydrophobic AWI can also cause non-uniform distribution of the particle orientations^16,20^, which hinders the reconstruction of 3D maps with isotropic resolution^21^. Five main sample preparation strategies to overcome AWI interactions have been pursued so far: (i) Specimen support films, (ii) chemical crosslinking, (iii) rapid freezing-plunging robots, (iv) specimen encapsulation, and (v) chemical additives, such as detergents or small proteins, which most likely act by coating the AWI with a continuous film.

Thin, amorphous carbon was among the first cryo-EM specimen support films used. Particles from the sample will adhere to the carbon, which will physically separate them from the AWI during grid preparation and may also serve to increase particle concentrations relative to unsupported grid preparation methods (e.g.^22–24^). However, amorphous carbon itself can also induce preferred orientation and may not be compatible with all samples. Therefore, additional types of specimen support films have been developed, including molecular graphene^25–28^, graphene oxide^29–32^, and affinity grids such as functionalised graphene oxide ^33^, antibody-coated grids ^34^, or streptavidin-affinity grids^35^. Although usage of an additional support layer has been found to be beneficial for many samples, the interaction of the sample with the support layer may also cause dissociation of the target macromolecular complex. Moreover, in-house production of these supports may be fiddly or technically difficult for some users and might introduce additional batch-to-batch variability.

Most support films can also be combined with chemical cross-linking of the sample. One of the most commonly used chemical crosslinkers in cryo-EM is glutaraldehyde, which stabilizes fragile macromolecular complexes and can improve the particle orientation distribution^36,37^. More recently, BS3, which is milder than glutaraldehyde, has gained more widespread application (e.g. ^38–44^). However, not all complexes can be stabilised by crosslinking, and the introduction of artefactual conformations by protein-protein crosslinking can be a concern.

Blot-free methods such as Spotiton^45^ or Chameleon^46^ allow extremely rapid plunge-freezing after sample deposition, which partially mitigates the harmful effects of the AWI because it reduces the time in which the sample particles can interact with it^47^. However, these robotic systems are costly to procure and maintain. Therefore, they are not accessible to many cryo-EM practitioners.

A variety of containers allowing the encapsulation of samples, thereby shielding them from the AWI, has been explored. This includes liposomes ^48^, protein nanocrates ^49^, and DNA origami structures ^50^. However, the relative complexity of these methods has so far precluded their wide-spread adoption in cryo-EM structure determination.

Finally, the usage of chemical additives to mitigate AWI interactions has been explored extensively. The effects of surfactants such as detergents (e.g. ^51–55^), amphipols (e.g. ^55,56^), or self-assembling monolayers ^57^ have been characterised in detail, and such additives have demonstrated their utility when applied to cryo-EM structure determination of challenging specimens (e.g. in^58–60^). However, a drawback of many detergents is that they substantially reduce the number of particles visualised in individual micrographs, therefore necessitating higher, and sometimes very high, sample concentrations ^51^. Additionally, some detergents also induce a loss of contrast in electron micrographs. More recently, denatured proteins^18^, or small protein domains interacting with the AWI^61,62^ have been explored as means towards preserving the structural integrity of the sample of interest. It has been surmised that both detergents and proteinaceous additives coat the AWI with a molecular film ^16,52,61^, thereby mitigating AWI interactions of the sample, though in some cases, it has been shown that particularly pronounced stabilising effects were caused by binding of the detergent to the target protein ^53^.

Here, we introduce cryo-tides, synthetic peptides of defined sequence, as a cost-effective and technically straightforward additive for cryo-EM grid preparation. Cryo-tides protect complex integrity and alleviate preferred particle orientations during cryo-EM grid preparation while preserving high image contrast and monodisperse sample distribution. We demonstrate the stabilising effects of our cryo-tides on the anaphase-promoting complex (APC/C) (reviewed in^63^), a fragile multi-subunit protein complex which hitherto required carbon support for particle preservation upon vitrification^22,64^. We have also applied the cryo-tides to structure determination of the small CDK11:cyclin L:SAP30BP complex ^65^, which exhibits strong preferred orientation.

## Results

### Cryo-tide design

In a previous study focusing on nucleosome samples, we observed that the addition of a small peptide derived from the cyclin B1 N-terminus greatly improved the yield of intact particles on cryo-EM grids^66^. This observation led us to hypothesise that this peptide might have protected the molecular complexes during vitrification. This peptide, which we refer to as the basic-hydrophobic (BH) cryo-tide (for *cryo*-EM pep*tide*), consists of 21 amino acids and features mainly nonpolar (43%) and basic (24%) residues. We subsequently rearranged this sequence to assess whether the effect we saw depended on the amino acid sequence, or merely on the amino acid composition of the peptide, thereby generating a scrambled basic-hydrophobic (SBH) cryo-tide. Moreover, to test the effect of positive and negative charges specifically, we made a charge swap version named the acidic-hydrophobic (AH) cryo-tide. Finally, we also used a 21 amino acid poly-lysine peptide (PL) to see if the protective effect on the sample was purely conveyed via the basic charges in the BH cryo-tide (**Figure 1**).

**Figure 1:**
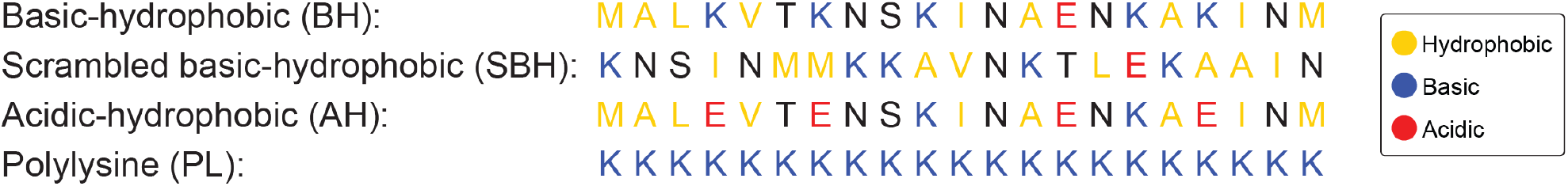
The sequences of the cryo-tides used in this paper. The peptide sequence are shown above with the residues coloured based on their chemical properties. Hydrophobic residues are marked in light-orange, basic residues in blue and acidic residues in red.

We first explored the effects of these four cryo-tides on the stability of the anaphase promoting complex/cyclosome (APC/C) during cryo-EM sample preparation. The APC/C is a 1.2 MDa complex formed by 19 subunits. It can be divided into three separate subdomains. The tetratricopeptide repeat (TPR) lobe is formed by two copies each of APC3, APC6, APC7, APC8, APC12 along with APC13 and APC16. The platform module is composed of APC1, APC4, APC5, and APC15, and the catalytic module is comprised of APC2, APC10 and APC11 (**Figure 2**). The homodimers of APC3, 6, 7, and 8 stack to form the TPR lobe. This region of the APC/C contains some inherent flexibility because this region can ‘breathe’ slightly or extend and contract. Because of the latter, the top of this lobe, where the APC7 homodimer is located, often presents weaker density in cryo-EM 3D reconstructions.

**Figure 2:**
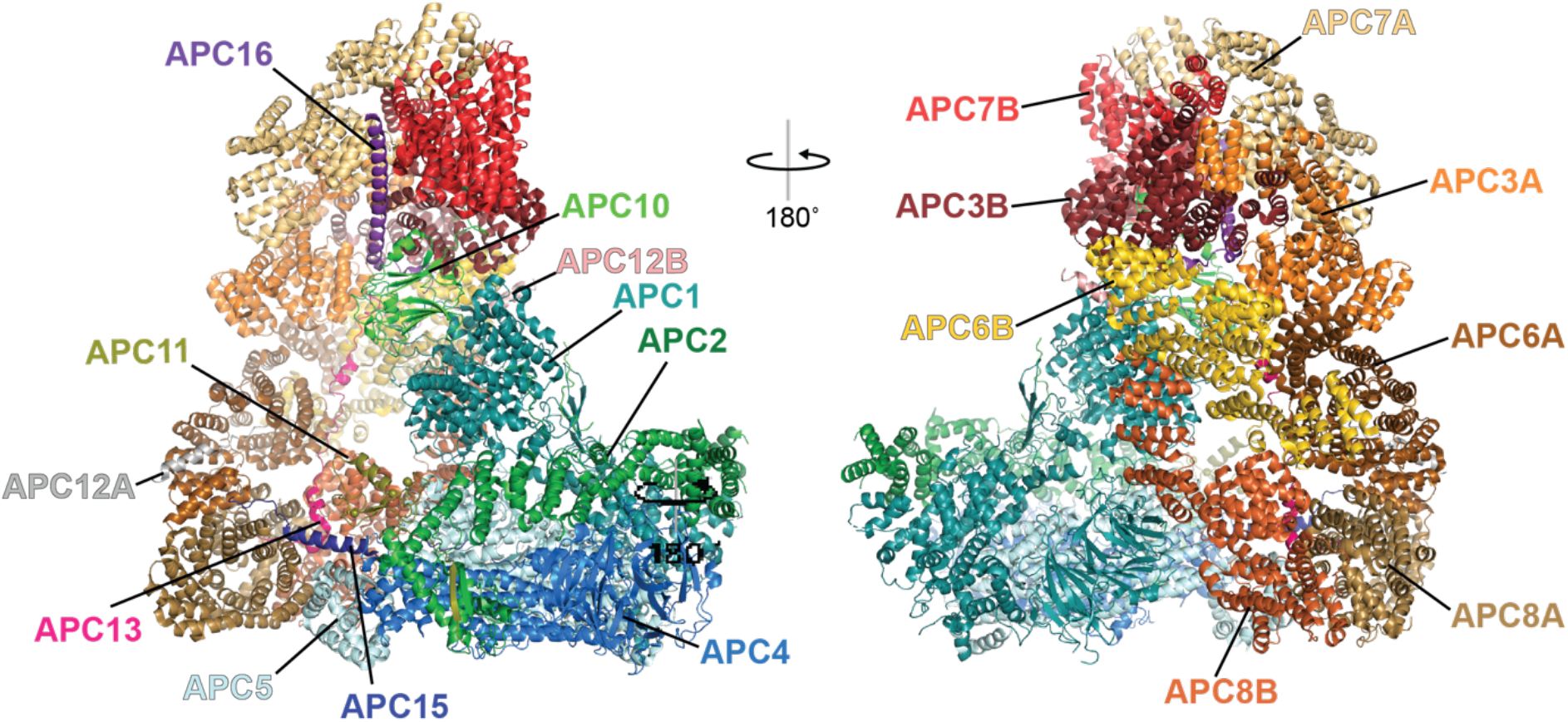
Overview of the structure of the APC/C. The structure of the APC/C with its 19 subunits individually coloured and labelled. PDB 5G05 was used to make this figure^70^.

The APC/C is an ideal test specimen for our cryo-tide system for several reasons. First, cryo-EM sample preparation of APC/C specimens typically requires the use of continuous carbon support of ∼50 Å thickness, which is applied to the grid prior to vitrification ^64,67–70^. Without this support film, the APC/C tends to fall apart. Therefore, using the APC/C, observing the effects of the cryo-tides on particle integrity and quality is straightforward and can be achieved by collection of single-particle cryo-EM data. Second, the complex is large and structured, facilitating particle picking on tomograms, which we used to assess particle coverage across the grid hole and the overall behaviour of the sample on the grid. We thus collected single particle micrographs and tomograms on the same grid for each sample, using a 200 kV Glacios cryogenic transmission electron microscope.

### Single Particle Analysis shows a higher number of intact particles in presence of the cryo-tide

Initial single particle image processing was performed in CryoSPARC ^71^. We picked particles using a combination of blob-, template- and Topaz-based ^72^ particle picking. After 2D classification, selected particle classes were subjected to *ab initio* reconstruction. The quality of these initial reconstructions is strikingly variable. As expected, given extensive prior studies of the complex, the control-APC/C sample falls apart in the absence of a carbon support. The *ab initio* class obtained contains the platform module only while the TPR lobe density is absent (**Figure 3A**), even though this class contained the most particles across the five datasets (**Table 1**). Conversely, *ab initio* reconstructions obtained from the BH, SBH and AH cryo-tide samples clearly show the complete APC/C. This can also be seen in the 2D classes, where the data obtained in the presence of these three cryo-tides contain significantly fewer views of broken APC/C particles when compared to the control sample (**Figure 3B, C and D**). In stark contrast, the PL peptide had an adverse effect on the APC/C sample. Each micrograph contained very few APC/C particles and those present were largely broken (**Figure 3E**). The PL peptide appeared to have an adverse effect not only on the protein within the sample but also on the quality of the electron micrographs obtained during data collection. Of the approximately 5500-6500 micrographs for each sample, approximately 50% were rejected during pre-processing in the PL cryo-tide dataset (**Table 1**). This was often due to the ice being devoid of particles, too thin, or too thick. The quality of the protein is also reflected in the percentage of broken vs. unbroken APC/C particles contained in the 2D classes obtained straight after blob picking. The intact APC/C 2D classes account for 25 % of the particles in the classification for the control sample, which is further decreased to 17 % in the PL peptide sample. However, when the BH, SBH and AH cryo-tides are present, the ratios swap, and 75-90 % of particles are contained in the 2D classes show intact APC/C views (**Table 1**). Overall, the BH, SBH and AH cryo-tides appear to be providing some level of protection, as shown by the increasing particle numbers and better particle integrity on the grids. Finally, although the number of broken particles is similar between the BH and the SBH cryo-tide, the AH cryo-tide grid presents about two-fold more broken particles, suggesting that the positive charges in the BH peptide are more beneficial than the acidic ones in the AH peptide.

**Figure 3:**
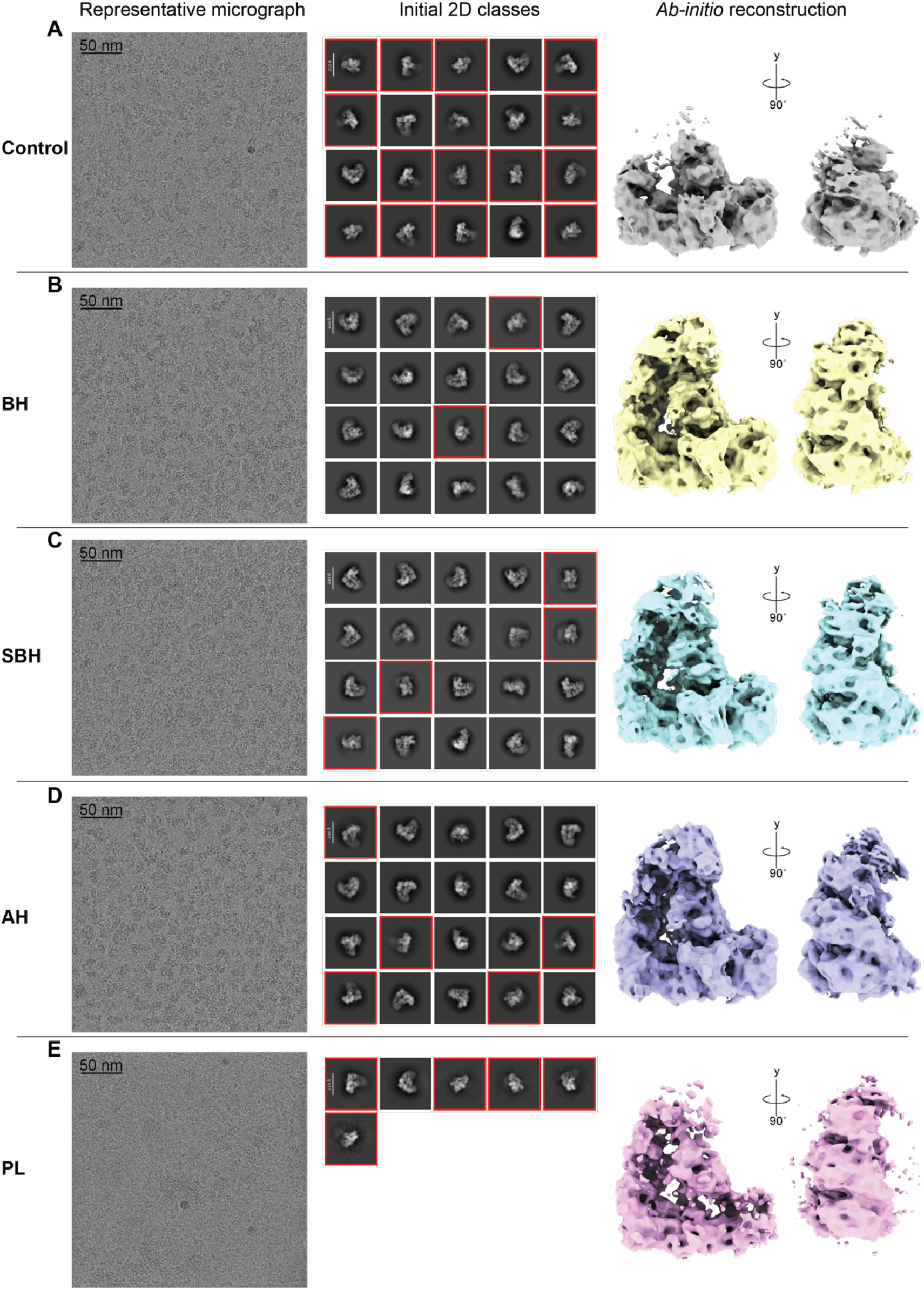
Initial processing of the APC/C samples. (**A-D**) For each sample a representative micrograph is shown on the left, the initial 2D classes selected after blob picking in the middle and the *ab initio* reconstruction gained after multiple rounds of picking to obtain the best classes on the right. Only the first 20 2D classes are shown, red boxes have been used to highlight classes of broken APC/C. After the blob picking all classes that contained well aligned protein particles were picked whether they were broken or not. The *ab initio* maps shown were opened and displayed at the same threshold values using ChimeraX^73^.

**Table 1:** Initial processing of the various APC/C datasets. For each sample the numbers of micrographs collected, and the numbers accepted after pre-processing are shown. The initial number of particles picked from each dataset by the cryoSPARC Live blob picking algorithm are also shown with, these particles were put into a 2D classification and a rough estimate of broken vs intact particles was calculated based on the numbers of particles that made up broken or intact APC/C views in the 2D classification. The final column shows the number of particles that make up the *ab initio* classes shown (**Figure 3A-D** rightmost panel). These were particles picked via blob-picking, template-based picking and Topaz based particle picking.

| **Sample** | **Micrographs collected** | **Micrographs accepted** | **Particles after CryoSPARC Live blob picking** | **Intact particles (%)** | **Broken particles (%)** | **Particles in *ab initio* reconstruction** |
| --- | --- | --- | --- | --- | --- | --- |
| Control | 5,500 | 5,022 | 203,962 | 24.3 | 75.7 | 403,199 |
| BH | 5,501 | 4,942 | 324,463 | 90.7 | 9.3 | 324,157 |
| SBH | 5,501 | 4,408 | 278,590 | 86.3 | 13.7 | 338,554 |
| AH | 6,506 | 4,029 | 265,378 | 75.9 | 24.1 | 282,966 |
| PL | 5,502 | 2,748 | 35,357 | 17.4 | 82.6 | 10,272 |

In conclusion, we observe that a hydrophobic and basic peptide increases the number of intact APC/C particles. This effect primarily depends on the amino acid composition rather than on the linear peptide sequence, and a mixed basic and hydrophobic peptide is slightly more protective than a mixed acidic and hydrophobic peptide. Finally, the hydrophobic component of the peptide is essential for its protective action.

### Different cryo-tides induce different particle orientation distributions

Particles assigned to the *ab initio* classes described in the preceding section were then used to perform a homogenous refinement using a single reference map, EMD-17751^74^. Therefore, all refinement outputs had a common reference frame and their viewing direction distribution heatmaps are comparable to each other (**Figure 4**). After homogeneous refinement, the control and PL samples do not show TPR lobe density, and the PL sample does not contain the catalytic module either, whereas the BH, SBH and AH cryo-tide maps show the intact APC/C. The viewing direction distribution heatmaps show that the particle orientation distributions in the BH and SBH samples are similar to one another, but different compared to the control sample. The sample containing the AH peptide shows the same main view, and in addition, a different set of secondary views, with prevalent new viewing directions graphed across the “bottom” half of the heatmap, as compared to the control, BH, and SBH samples (**Figure 4B, C, and D**). This indicates that the cryo-tides are not only capable of protecting and enriching APC/C particles, but that they also affect the orientations of the complex on grid. Under the assumption that the orientation of the particles on the grid depends on the local surface properties of the particles themselves and the propensity of these local particle surface areas to interact with the interface layer^16^, this result is to be expected. Therefore, in addition to preserving complex integrity, cryo-tides may also help tackling issues with orientation bias. Compared to the APC/C reconstructions in the presence of BH, SBH, and AH cryo-tides, APC/C imaged in the presence of the PL cryo-tide exhibits strong preferred orientation, with a few highly populated views and very little coverage of the remainder of the Euler angles. The Euler angle distribution maps are useful to identify the most dominant views of the APC/C (**Figure 4A-E rightmost panel**).

**Figure 4:**
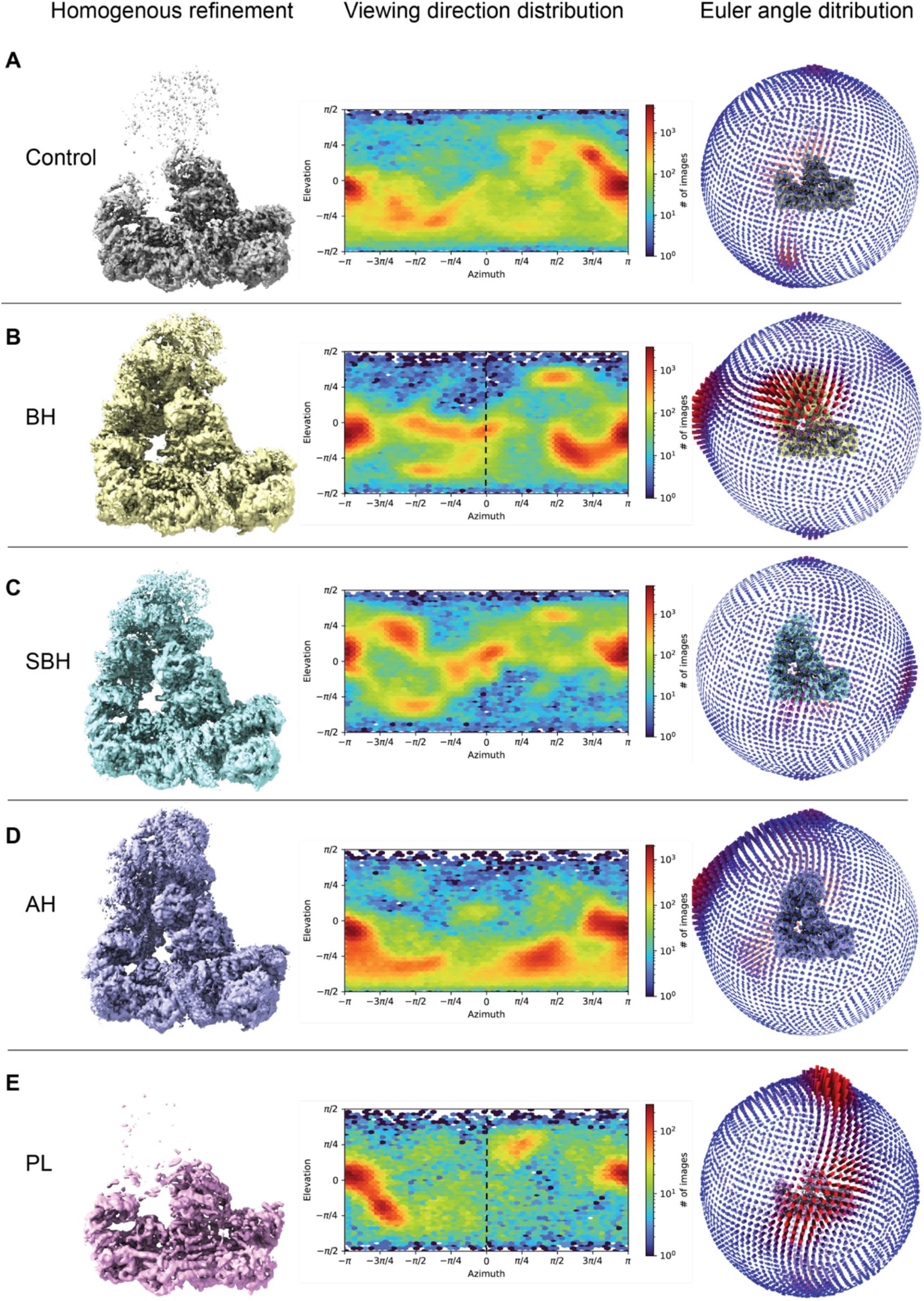
Orientation distributions of the APC/C samples. For each sample, a heatmap of the viewing direction distribution (middle) of the particles included in the homogenous refinement (cryo-EM map on the left) is shown. To make all heat maps directly comparable, some of them had to be translated by 180° on the azimuth axis and mirrored across 0° elevation to account for the fact that imaging a particle under projection angles differing by 180° (e.g. because a grid was loaded “upside down” or because the particles occupy the opposite AWI) generates projections related by mirror symmetry. Heatmaps transformed in this way are marked by dashed black lines at 0 azimuth. The Euler angle distribution map for each sample is shown on the right. Longer red lines represent more highly represented views.

We did not further process the control and PL datasets after this stage because they did not yield high resolution reconstructions of the complete APC/C particle. The BH, SBH and AH samples, each of them providing a high-quality APC/C map, were subjected to more extensive processing in RELION^75,76^. After extensive processing, including 3D classification, Bayesian polishing, and 3D auto-refinement, the BH, SBH and AH datasets yielded reconstructions of the APC/C at 3.3 Å, 3.4 Å, and 3.4 Å global resolution, respectively (**Figure 5A-C**). For all these maps, the core of the APC/C, comprising the platform and lower half of the TPR lobe, contains the regions of best resolution. The maps become more fragmented at the top of the TPR lobe and at APC2, APC11, and APC4 (**Figure 2** and **Figure 5**). The latter regions are commonly the least rigid parts of the APC/C. Therefore, lower quality density in these regions was expected. Even though all three maps have almost the same reported resolution, visual inspection indicates that the BH and SBH maps are of better quality overall than the AH map. At the same threshold values, the AH map exhibits weaker density for the TPR lobe and at the edges of the APC2/APC4 subunits at the periphery of the complex (**Figure 5C**). This is illustrated in the APC16 helix which sits in the centre of the TPR lobe. Both the BH and SBH samples have stronger density here and in the surrounding regions than the AH sample reconstruction (**Figure 5A-C**).

**Figure 5:**
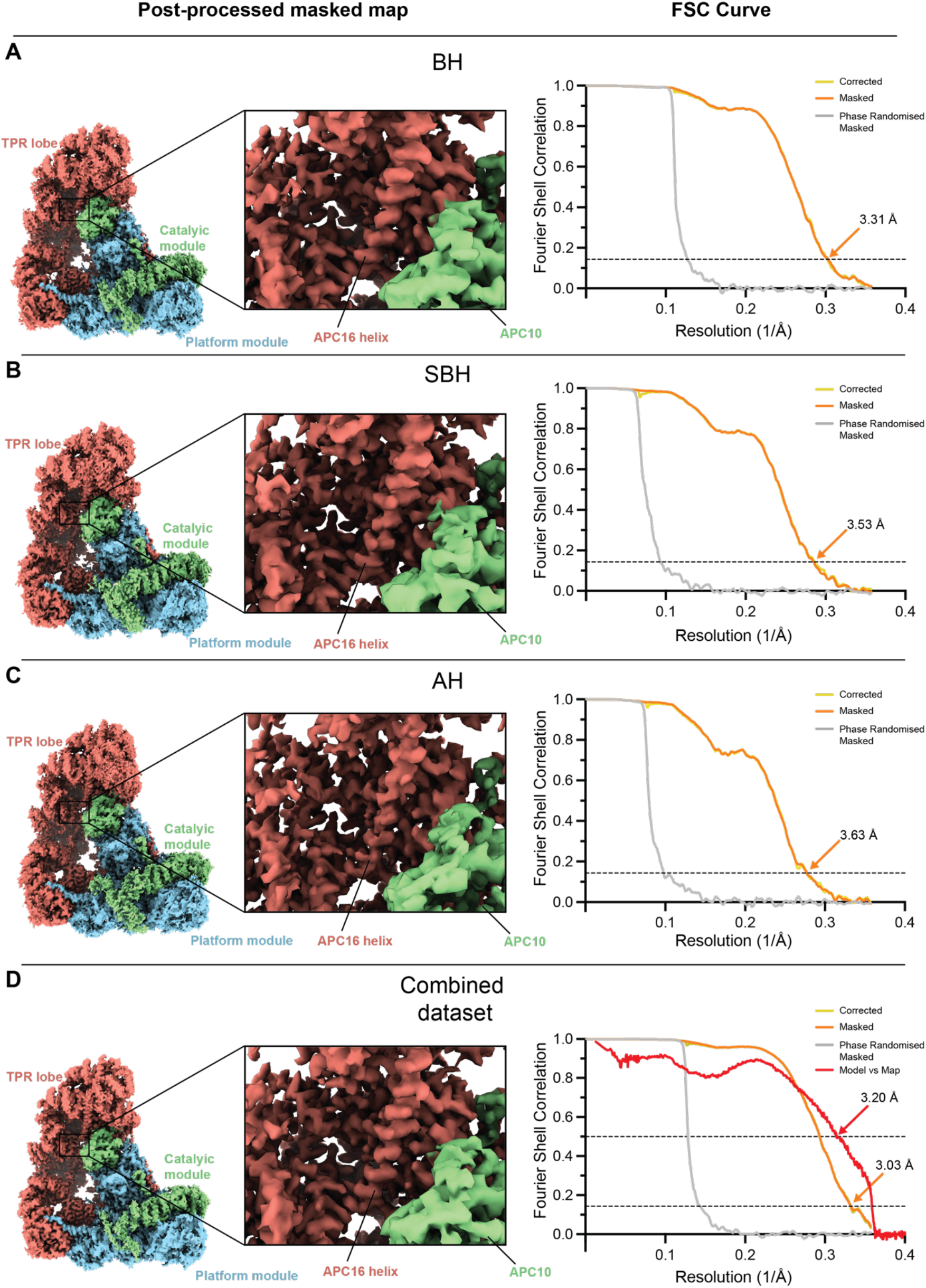
Final postprocessed maps from cryo-tide datasets. (**A-D**) The final postprocessed map of the various datasets. The final post-processed maps are shown on the left. The TPR lobe for each map is coloured in red, the catalytic module in green and the platform module in blue. The middle of each section shows a zoomed in picture of the APC16 helix and the right-hand side shows the FSC curves of the corrected (yellow), masked (orange), phase randomised (grey) and model vs. map (red) maps.

In summary, different cryo-tides featuring a combination of hydrophobic and charged residues protect particle integrity and induce different particle orientation distributions.

### Different datasets of the same sample with different cryo-tides can be pooled to generate more isotropic reconstructions

To investigate whether combining the BH, SBH, and AH datasets with their non-identical orientation distributions would further increase the quality and resolution of the APC/C cryo-EM reconstruction, we pooled the final particle sets and ran 3D auto-refinement on this combined dataset. The SBH and AH samples contained ∼64,000 and ∼69,000 particles, respectively, while the final reconstruction of the BH sample contained ∼93,000 particles. The map computed from the pooled datasets was further improved, with a global resolution of 3.0 Å. This map exhibited the best density overall, with a slightly improved upper TPR lobe (**Figure 5D**).

We then compared the final viewing direction distributions and the directional FSC ^21^ to visualise the combined orientation distribution and to investigate whether combining of the datasets had reduced the anisotropy of the resulting map (**Figure 6**). The viewing distribution plots of the three fully processed datasets showed varying degrees of preferred orientation, with the SBH and AH datasets showing more pronounced orientation bias and a reduced 3D FSC sphericity (0.84 and 0.87, respectively; **Figure 6A-C**). As expected, the orientation distribution of the combined dataset is the sum of the orientation distributions of the three input datasets and has a much greater apparent coverage of views (**Figure 6D**). The combined dataset exhibits the highest sphericity value (0.96) among all reconstructions in 3D FSC analysis, further improved beyond the BH dataset (at 0.93) (**Figure 6A-D**).

**Figure 6:**
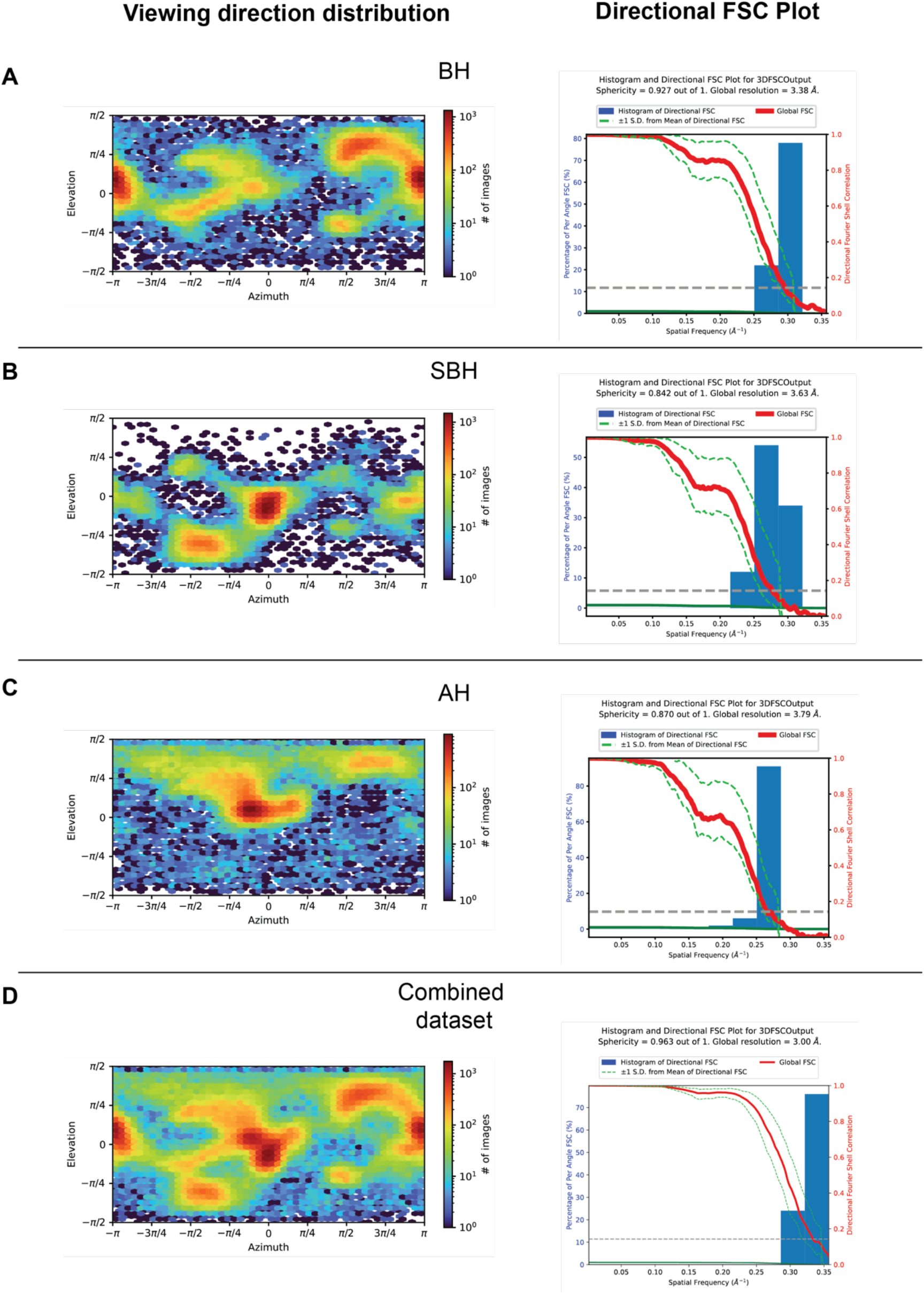
Particle orientation information from the final 3D refinements for each dataset. (**A-D**) For each sample the viewing direction distribution heatmaps generated by CryoSPARC are shown on the left and the directional 3DFSC histograms are shown on the right. The directional 3DFSC histograms were generated using 3DFSC software ^21^.

To conclude, the cryo-tides not only protect the sample from denaturation but also modify the particle orientations present. Therefore, by combining datasets collected in the presence of different cryo-tides, it is possible to further improve the resolution and map quality over individual datasets. This approach provides a fast and cost-effective way of reducing orientation bias during sample preparation.

### Cryo-tides influence the thickness of particle layers through the ice

To obtain further insights into the effects of the cryo-tides on the distributions of the APC/C particles within the ice layer, we subsequently analysed the cryo-tide APC/C samples using cryo-electron tomography (cryo-ET). To do this, we collected tomograms on the same grids that had been used for the single-particle analysis.

Once reconstructed, the tomograms looked different to one another by visual inspection. When looking at a central slice in the tomograms, it is possible to see individual APC/C particles (**Figure 7**). Visual inspection readily reveals that the cryo-tide-containing samples, with the exception of the PL sample, manifest higher numbers of distinct individual particles in the tomogram when compared to the control sample (**Figure 7A-D**). The control sample also contains a grainier background, and it is thus more difficult to pick out individual, distinct particles (**Figure 7A**). This could be due to the large number of broken protein particles that were detected in single particle analysis, which matches with the smaller grains not seen in the BH, SBH, and AH cryo-tide samples. As expected after the single particle analysis, the PL sample contained very few identifiable protein particles and those present were largely found at the edge of the holes. These PL sample tomograms also show a large amount of aggregation or empty ice (**Figure 7E**).

**Figure 7:**
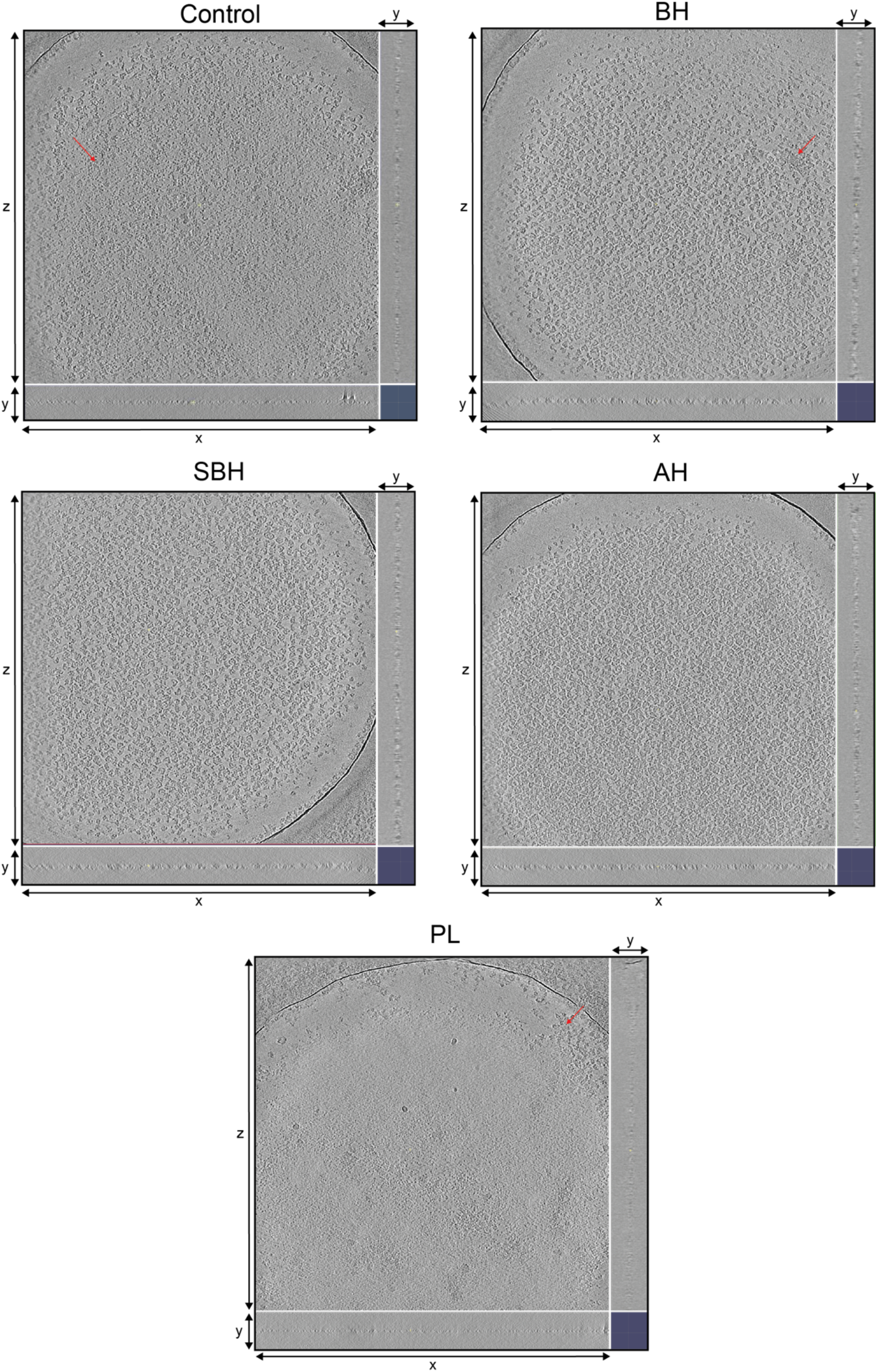
Images showing a central slice of each sample. A central slice from a representative tomogram of each sample. The large square plane shows the view of the x,z axes, underneath are the x, y axes and to the right the y, z axes. Some examples of aggregation are shown with a red arrow, not all potential aggregation is highlighted. Images made using IMOD ^77^.

The particles in the tomograms were picked, and their positions were plotted onto a graph. For each sample, a 200-pixel range along the x axis was selected, particles were picked inside the volume defined by this 200-pixel segment along the x axis and the full range along the y- and z-axes contained in the tomogram, and then the particle locations were plotted on a graph by collapsing the x-axis (**Figure 8A**). This analysis revealed that the control APC/C sample contains fewer particles in the test volume when compared to the BH, AH and SBH samples. In the control sample, these particles are distributed in a thin layer through the ice. The sample layer only thickens near the carbon edges. In contrast, the BH, SBH and AH samples all contain a thicker layer of APC/C particles throughout the ice (**Figure 8C**). This, combined with the fact that the particles are 70-90% intact in the initial processing of the cryo-tide samples and only ∼25 % intact in the control sample (**Table 1**), suggests that the cryo-tides are providing a protective layer for the protein complex, which prevents it from adhering to and falling apart at the AWI. Similarly, the number of particles picked from the tomograms was greater with the BH, SBH and AH samples when compared to the control sample (**Figure 8B**). Finally, the PL sample shows very few particles across the centre of holes with most of the particles being present at the carbon edge of the hole across the whole tomogram. The average number of picked particles within the 200 px tomogram slices was drastically lower than the control sample (**Figure 8B and C**).

**Figure 8:**
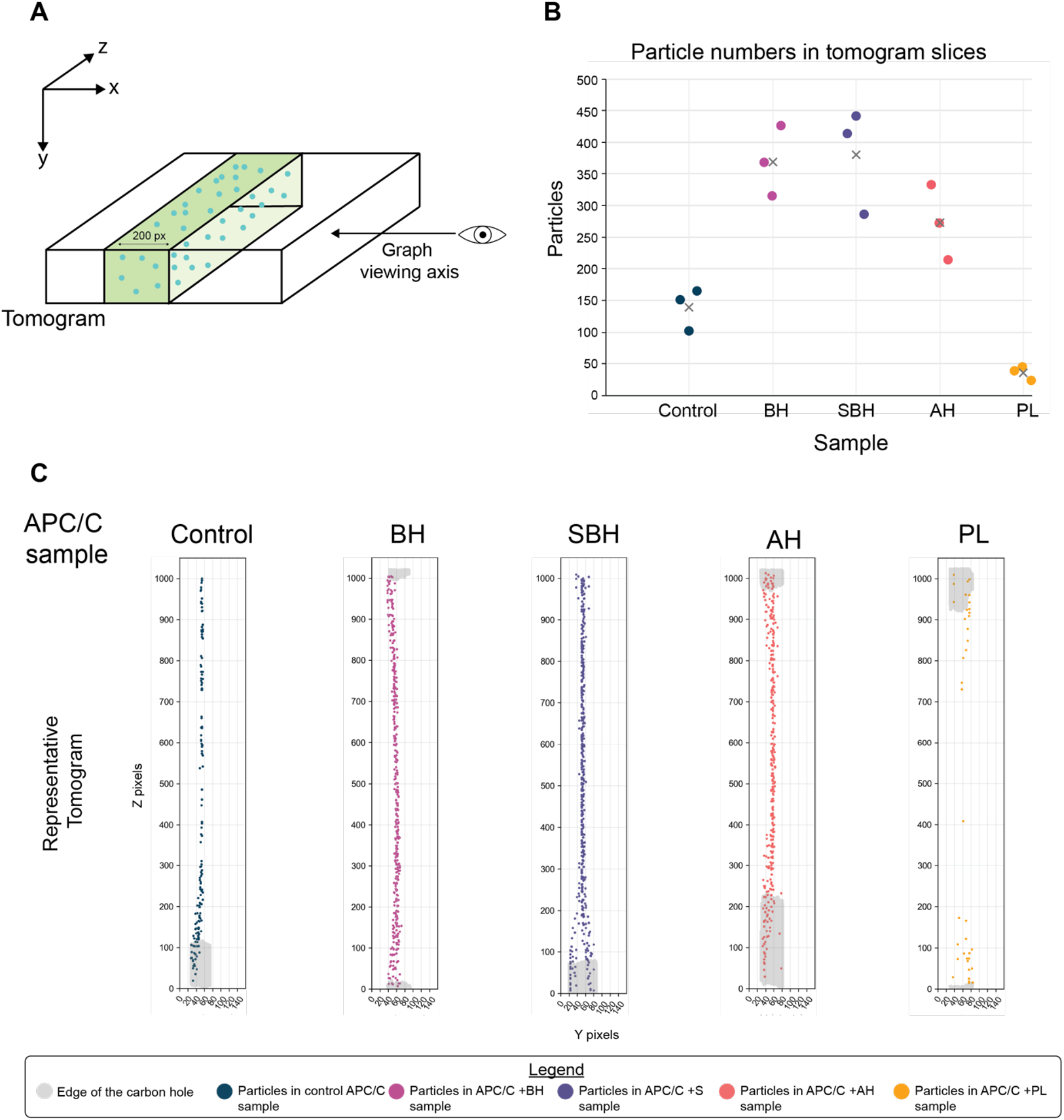
Particles picked in tomograms. (**A**) A schematic showing how the boundaries were defined and the labelling of the axis to aid with understanding of the plots below in section. The effect of the missing wedge is not considered in this depiction. (**B**) A graph showing the total number of particles picked in three 200 px tomogram slice for each sample. The final number picked in each slice is represented by a circle marker and the mean number of particles for each sample is shown by a grey cross. (**C**) Plots showing the picked particles in the ice of the tomograms. For both **B** and **C** the control sample particles are shown in dark blue, the BH in pink, SBH in purple, AH in orange, and PL in yellow. The carbon edge is shown in transparent grey.

In conclusion, the cryo-tides increase the thickness of the particle layer through the ice, thereby increasing the number of particles. By preventing and partially replacing particle interactions with the AWI, the cryo-tides are likely to exert the effect on particle orientation distributions observed by single particle analysis.

### The cryo-tide is instrumental for determining the cryo-EM structure of CDK11:cyclin L:SAP30BP complex

In addition to enabling structure determination of the APC/C in the absence of carbon support, the cryo-tide approach also facilitated structure determination of the CDK11:cyclin L:SAP30BP complex. During sample optimisation for our published structure of this complex ^65^, we initially collected 7,620 electron micrograph movies (7,231 accepted for further processing) from specimens prepared using conventional methods, which resulted in a 3.5 Å-resolution map with streaky appearance, a sign of anisotropic resolution (**Figure 9A, B**, **Supplementary Fig. 8**). Map appearance and resolution (3.4 Å) were improved after inclusion of the BH cryo-tide during sample preparation (**Figure 9C, D**, **Supplementary Fig. 8B**), notably from only 5,087 movies (4,161 accepted for further processing) and fewer particles (259,564 particles, as compared to 418,797 for the sample prepared without cryo-tide, see Methods and **Supplementary Fig. 8**). Further analysis indicated that the BH cryo-tide leads to a broader particle orientation distribution, as judged by orientation distribution plots obtained from cryoSPARC ^71^ homogeneous refinement (**Figure 9E, F**), consistent with the appearance of additional high-quality 2D class averages in 2D classification (**Figure 9G, H**). While the resolution histogram obtained from 3D FSC analysis ^21^ of the specimen prepared using the CDK11-containing complex alone revealed a strong lower-resolution peak at around 5-6 Å resolution, indicative of poorly resolved viewing directions, this low-resolution peak was strongly reduced in the presence of the cryo-tide (**Figure 9I, J**). Furthermore, the conical Fourier area ratio (cFAR)^78^ for the reconstruction in the presence of the BH cryo-tide was improved as well, demonstrating a quantifiable improvement of map anisotropy. We thus used the BH cryo-tide for determination of our published 2.3 Å-resolution structure of the CDK11:cyclin L:SAP30BP complex. ^65^

**Figure 9:**
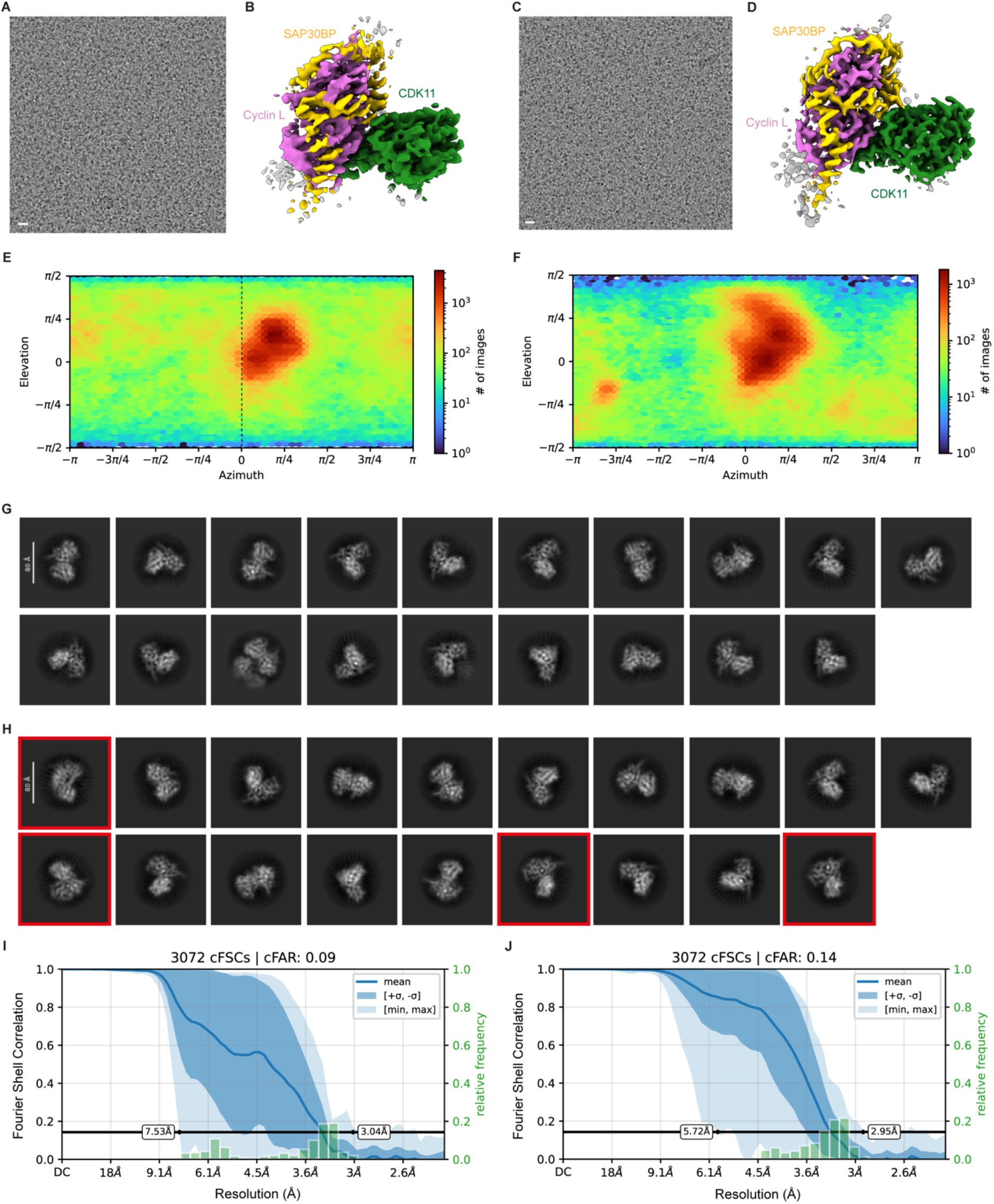
Cryo-tide-supported structure determination of the CDK11:cyclin L:SAP30BP complex. (**A**) Sample micrograph of CDK11:cyclin L:SAP30BP particles in vitreous ice. Scale bar: 100 Å. (**B**) 3D reconstruction of the CDK11:cyclin L:SAP30BP complex determined in the absence of cryo-tide. (**C**) Sample micrograph of CDK11:cyclin L:SAP30BP particles in vitreous ice in the presence of cryo-tide. Scale bar: 100 Å. (**D**) 3D reconstruction of the CDK11:cyclin L:SAP30BP complex determined in the presence of cryo-tide. (**E**, **F**) Orientation distribution plots from cryoSPARC homogeneous refinement for the CDK11:cyclin L:SAP30BP complex in the absence (E) and presence (F) of cryo-tide. (**G**, **H**) 2D class averages obtained from CDK11:cyclin L:SAP30BP samples in the absence (G) and presence (H) of cryo-tide. Classes in h without equivalents in G are highlighted. (**I**, **J**) Analysis of anisotropy of the CDK11:cyclin L:SAP30BP cryo-EM reconstructions using directional FSC and cFAR analysis ^21,78^.

## Discussion

We have shown that cryo-tides are a powerful tool to improve the quality of cryo-EM samples and data. We found that the inclusion of the cryo-tides reduces the amount of complex breakage for a fragile sample during sample preparation. Given the extensive evidence indicating that protein complexes break apart at the AWI, these peptides likely exert their effects by preventing the vulnerable regions of the protein from directly interacting with the highly hydrophobic AWI.

Furthermore, we have shown that the cryo-tides can influence particle orientations within the ice. Therefore, they can be used to induce different particle orientations, which can lead to an improvement in resolution and map quality. Due to their different physico-chemical properties, different cryo-tides can induce different particle orientations. Therefore, combining different cryo-tide datasets can be used to improve the coverage of viewing directions, reduce map anisotropy, and improve overall data quality when compared to using one cryo-tide alone. The BH and SBH cryo-tides have the same charge and number of basic and hydrophobic residues whereas the AH sample has a charge reversal. The differences between the particle orientations between the BH/SBH and AH cryo-tides must originate from the physical and chemical properties of the peptide, most likely their different charge, given that the BH and SBH samples contain largely similar particle orientation distributions whereas the orientation distribution in the AH sample is substantially different. This suggests that the interaction of the charged residues with the sample and/or the AWI influences how the protein can lie in the ice. We note that this observation agrees with prior analysis of ionic detergents in which positively and negatively ionic detergents were found to induce different sets of particle orientations ^52,55^. It is important to note that the data quality of the PL sample was poor, the initial 3D refinements yielded maps of worse quality compared to the control sample, and very few particles could be picked. This shows that it is not only the charge (basic/acidic residues) of the peptide that is important, but that the hydrophobic residues must also be playing a role in the protection.

Finally, we were able to show that there were more visible intact, undamaged, particles present on the grids made from samples containing cryo-tides, and that these particles occupy a thicker layer when compared to the control sample, especially when comparing the centre of the holes. This could be due to the peptide preventing the denaturation of the particles, and therefore, the particles prevent the ice film from “collapsing” to a very thin layer that contains only denatured debris.

To conclude, the use of cryo-tides in cryo-EM sample preparation is a cheap, time efficient, and simple tool to use during cryo-EM sample vitrification. These cryo-tides represent a two-pronged approach towards improving the cryo-EM sample and thereby the quality of the final map: by protein protection during vitrification and via more varied particle orientations obtained, thereby reducing map anisotropy. We have observed that cryo-tides have helped with cryo-EM grid preparation of several samples including nucleosomes^66^, SIN3B complexes^79^ and, as shown here, the APC/C and CDK11:cyclinL:SAP30BP ^65^. We therefore envision that our cryo-tides can support structural analysis of a wide range of samples. Moreover, our work can be used as a base for the design of additional, more diverse cryo-tides that may be advantageous for an even wider range of samples.

## Methods

### Protein expression and purification

The APC/C was expressed in insect cells using two separate baculoviruses which were created in the lab of David Barford^80^. The first virus contained APC/C subunits: APC5, APC8, APC10, APC13, APC15, APC2 and APC4-strepII and the second virus: APC1, APC3, APC6, APC7, APC11, APC12 and APC16. High5 cells (*Trichoplusia ni*; Thermo Fisher Scientific, cat. no. B85502) at a cell density of 2X10^6^ were infected with the baculoviruses and incubated for 72 hours at 130 rpm and 27 °C.

The APC/C cell pellets were thawed on ice in wash buffer [50 mM HEPES pH 8.3, 250 mM NaCl, 5 % glycerol, 2 mM DTT, 1 mM EDTA, 2 mM Benzamidine] supplemented with 0.1 mM PMSF, 5 U/mL Benzonase and 1 EDTA-free protease inhibitor tablet (Roche). The cells were lysed by sonication on ice and then centrifuged at 48,000 x g for 1 hour at 4 °C. The supernatant was bound to 3 x 5ml StrepTactin Superflow Plus cartridges (QIAgen) using a 1.5 mL/min flow rate and the columns were washed extensively with wash buffer at pH 8.0. The bound APC/C was eluted from the column using wash buffer pH 8.0 supplemented with 2.5 mM desthiobiotin (IBA-Lifesciences). The APC/C was incubated overnight at 4 °C with tobacco etch virus (TEV) protease overnight. The next morning the sample was diluted 2-fold with saltless buffer A [20 mM HEPES pH 8.0, 125 mM NaCl, 5% glycerol, 2 mM DTT, 1 mm EDTA]. The sample was then further purified using anion exchange chromatography by loading onto a 6 mL Resource Q column (GE Healthcare) with an elution gradient of 0-100% buffer B [20 mM HEPES pH 8.0, 1 M NaCl, 5% glycerol, 2 mM DTT, 1 mm EDTA] over 125 mL. The APC/C was concentrated and buffer exchanged into a final buffer [20 mM HEPES pH 8.0, 150 mM NaCl, 0.5 mM TCEP] and then centrifuged (Optima TLX Ultracentrifuge) at 40,000 rpm, 4 °C for 30 minutes.

For preparation of CDK11:cyclin L:SAP30BP, we used the baculoviral protein expression constructs described previously^65^: Twin-strep-tagged CDK11^p1^^10^ residues 357-795, His_6-_tagged cyclin L2 residues 1-319, and MBP-tagged SAP30BP. The complex was expressed in High5 cells after two rounds of virus amplification in Sf9 cells (*Spodoptera frugiperda;* Thermo Fisher Scientific, cat. no. 11496015). Cells were harvested and frozen at -80 °C after 72 h of incubation post infection.

The harvested cells were then thawed in purification buffer (250 mM KCl, 2 mM MgCl_2,_ 40 mM HEPES-KOH pH 7.9, 5 mM β-mercaptoethanol, 10% (v/v) glycerol) supplemented with protease inhibitors and DNase I. Cells were lysed by successive cycles of freeze-thaw, and cell debris were pelleted by centrifugation at 18,000 rpm (approx. 26,600 × g) for 30 min at 4 °C in a JA-25.50 rotor (Beckman Coulter). The CDK11:cyclin L:SAP30BP complex was subsequently purified by sequential Strep-Tactin affinity chromatography (elution by 10 mM desthiobiotin in purification buffer), cleavage of affinity tags by TEV protease, and size exclusion chromatography using a Superdex 200 Increase 10/300 GL column (Cytiva). The SEC and final storage buffer comprised 200 mM KCl, 2 mM MgCl_2,_ 20 mM HEPES-KOH pH 7.9, 5 mM β-mercaptoethanol, 5% (v/v) glycerol. The peak elution was pooled, concentrated to approximately 2 mg/mL, and snap frozen in liquid N_2 b_efore storage at -80 °C.

### Cryo-tide preparation

The unmodified basic-hydrophobic peptide [MALKVTKNSKINAENKAKINM], the scrambled basic-hydrophobic peptide [KNSINMMKKAVNKTLEKAAIN] and the polylysine peptide [KKKKKKKKKKKKKKKKKKKKK] were ordered from GenScript. The unmodified acidic-hydrophobic peptide [MALEVTENSEINAENEAEINM] was ordered from BioSynth.

### Grid preparation

The peptides were gently resuspended in cryobuffer [20 mM HEPES pH 8.0, 150 mM NaCl, 0.5 mM TCEP] to a final concentration of 2 mM at room temperature. For the APC/C samples, peptide at the final concentration of 200 µM or cryobuffer [20 mM HEPES pH 8.0, 150 mM NaCl, 0.5 mM TCEP] was added to 1 µM APC/C. QuantiFoil R1.2/1.3 Cu 300 grids were glow discharged using 15 mA for 1 minute on each side using the Easiglow (Pelco). 2 µL of sample was applied to the grid and blotted for 5 seconds using blot force 0 at 4 °C, 100 % humidity. The grids were frozen in liquid ethane using a Vitrobot Mark IV (Thermo Fisher Scientific).

Cryo-EM specimens of the CDK11:cyclin L:SAP30BP complex were prepared on UltrAuFoil R1.2/1.3 holey gold grids (Quantifoil Microtools). For grid preparation, the protein complex was diluted to 0.4 mg/mL in 200 mM KCl, 2 mM MgCl_2,_ 40 mM HEPES-KOH pH 7.9, and 1 mM AMP-PNP (Sigma-Aldrich), with and without 200 µM of BH cryo-tide (sequence MALKVTKNSKINAENKAKINM). 4 µL-samples were applied to plasma cleaned grids (Tergeo-EM plasma cleaner, PIE Scientific), mounted in a Vitrobot Mark IV (Thermo Fisher Scientific; operated at 5°C and 100 % humidity), blotted for 1, 1.5 or 2 sec, and plunged into liquid ethane cooled by liquid N_2._

### Cryo-EM Data Collection

For the single particle cryo-EM data collection of the APC/C with and without the various peptides, 5500 EER micrograph images were collected for each of the datasets on our in-house Glacios cryo-TEM 200 kV microscope. This was equipped with a Falcon4i detector, collecting in counting mode and with a physical pixel size of 0.7 Å per pixel and a 10 eV energy filter slit. A magnification of 165 kV was used with an electron dose of 60 e^-^/Å^2^ and a defocus range of -0.6 to -1.6 µm. A summary of APC/C data collection and processing statistics is provided in Supplementary Table 1.

Grids containing CDK11:cyclin L:SAP30BP complex were screened on a Glacios cryo-transmission electron microscope operated at 200kV acceleration voltage and equipped with a Falcon 4i direct electron detector. Data were acquired using the EPU software (Thermo Fisher Scientific) in EER format at a pixel size of 0.5675 Å/pixel and a total exposure of 60 electrons/Å^2^. A summary of CDK11:cyclin L:SAP30BP data collection and processing statistics is provided in Supplementary Table 2.

### Cryo-EM Data Processing

#### APC/C datasets

The micrographs were pre-processed on-the-fly using cryoSPARC Live, the micrographs were motion corrected and the initial 2D particle blob picking was carried out here. The data was then further processed in cryoSPARC with 2D classification followed by template particle picking and iterative Topaz training, Topaz particle picking, and 2D classification. The final selected particles were input into *ab initio* reconstructions with 3 classes. The selected *ab initio* class was input into a homogenous in cryoSPARC with cryo-EM map EM map EMD-17751^74^ as the reference, this was only to put the five maps into a common reference frame to compare particle orientations at this point. The particles making up the *ab initio* were converted into RELION format using csparc2star^81^. The particles were extracted in relion, then refined using the selected *ab initio* class as a reference, the particles were re-extracted and refined again. The SBH data was classified into 3 classes, class 2 selected and then went through 3D refinement, Bayesian polishing, CTF refinement and 3D refinement again before being classified with a mask for the TPR lobe. The selected particles were 3D refinement and post processed. The BH and AH samples went through Bayesian polishing, CTF refinement and 3D refinement twice before being classified for the best TPR lobe density. Again, the selected classes were refined and Post Processed to give the final reconstructions. The particles that made up the final 3D refinements for each dataset were refined with the BH final map low pass filtered as the starting model to put all the particles into a common reference frame. They were then refined together using the BH map as a reference, CTF refined and then a final 3D refinement carried out. This was post processed to give the final sharpened map (Supplementary Figs. 2-6).

#### CDK11:cyclinL:SAP30BP

CDK11:cyclinL:SAP30BP complex data were processed on-the-fly using cryoSPARC live ^71^, including motion correction, particle picking, and 2D classification. After the end of the data collection, all particles were re-extracted from the micrographs with re-centering, 2D classified again, and then subjected to homogeneous refinement. Based on this homogeneous refinement, particles were subdivided into two clusters based on their per-particle scale factors, and the particles contained in the higher cluster were used for the final homogeneous refinement (**Supplementary Fig. 8A, B**). The resulting reconstructions were resolved to approximately 3.5 Å (conventional grid preparation) and 3.4 Å (with cryo-tide).

### Coordinate refinement of the APC/C molecular model

Atomic coordinates from APC/C apo (PDB-ID: 8PKP)^74^ were fitted in the APC/C control map with Chimera. The fitted model was refined manually using COOT^82^ and real-space refinement was performed in PHENIX^83^. A summary of model refinement and validation statistics is provided in Supplementary Table 3.

### Cryo-ET Data Collection and Processing

For tilt series EER images were collected on the in-house Glacios 200 kV microscope equipped with a Falcon4i detector collecting in counting mode and collected with a 20 eV energy filter. Images were collected at 49 kV magnification with a 2.49 Å pixel size. The tilt series was collected using a -60° to 60° tilt scheme with an increment of 3° between each image. All tilt series images were collected at -7.00 µm defocus. The frames and tilt series were aligned and then the tomogram reconstructed using Etomo with a binning factor of 4 (ref. ^77^). An example of a reconstructed tomogram can be seen in Supplementary Movie 1. The particles were picked using Dynamo^84^.

## Data availability

The structural coordinates for the combined cryo-tide dataset has been deposited in the Protein Data Bank (https://www.rcsb.org/) with the accession number: PDB-ID 34MA. The EM data were deposited in the Electron Microscopy Data Bank (https://www.ebi.ac.uk/emdb/) with the accession codes: <u>EMD-100005 (BH dataset)</u>, <u>EMD-100028 (SBH dataset), EMD-100007 (AH</u> <u>dataset) and EMD-100029 (combined data)</u>. The wwPDBs logins are: The cryo-EM maps of the cryo-tide and control cryo-EM reconstructions of the CDK11:cyclin L:SAP30BP complex have been deposited in the Electron Microscopy Data Resource (EMDB) with accession codes EMD-59982 and EMD-59992, respectively.

## Author contributions

RY performed protein purification, cryo-EM grid preparation, data collection and processing for the APC/C samples, and wrote the manuscript with the help of CA and BJG. MJ helped with the latter data collection. BJG and AJSM provided the data for the CDK11 complex and contributed to the manuscript. CA coordinated the project, designed experiments, and helped with writing the manuscript.

## Acknowledgments

We would like to thank the ICR cryo-EM facility, in particular to Matthew Jessop for his help with cryo-EM data collection. We would like to acknowledge Jing Yang and Ziguo Zhang from David Barford’s laboratory for helping with the APC/C baculovirus generation. CA is funded by a Wellcome Career Development Award (302297/Z/23/Z). RY and MJ were funded by The Institute Of Cancer Research. BJG was supported by a career development fellowship from the Medical Research Council of the UK (grant number MR/V009354/1) and AJSM was funded by an ICR PhD studentship.

## Supplementary Materials

**Supplementary Figure 10:**
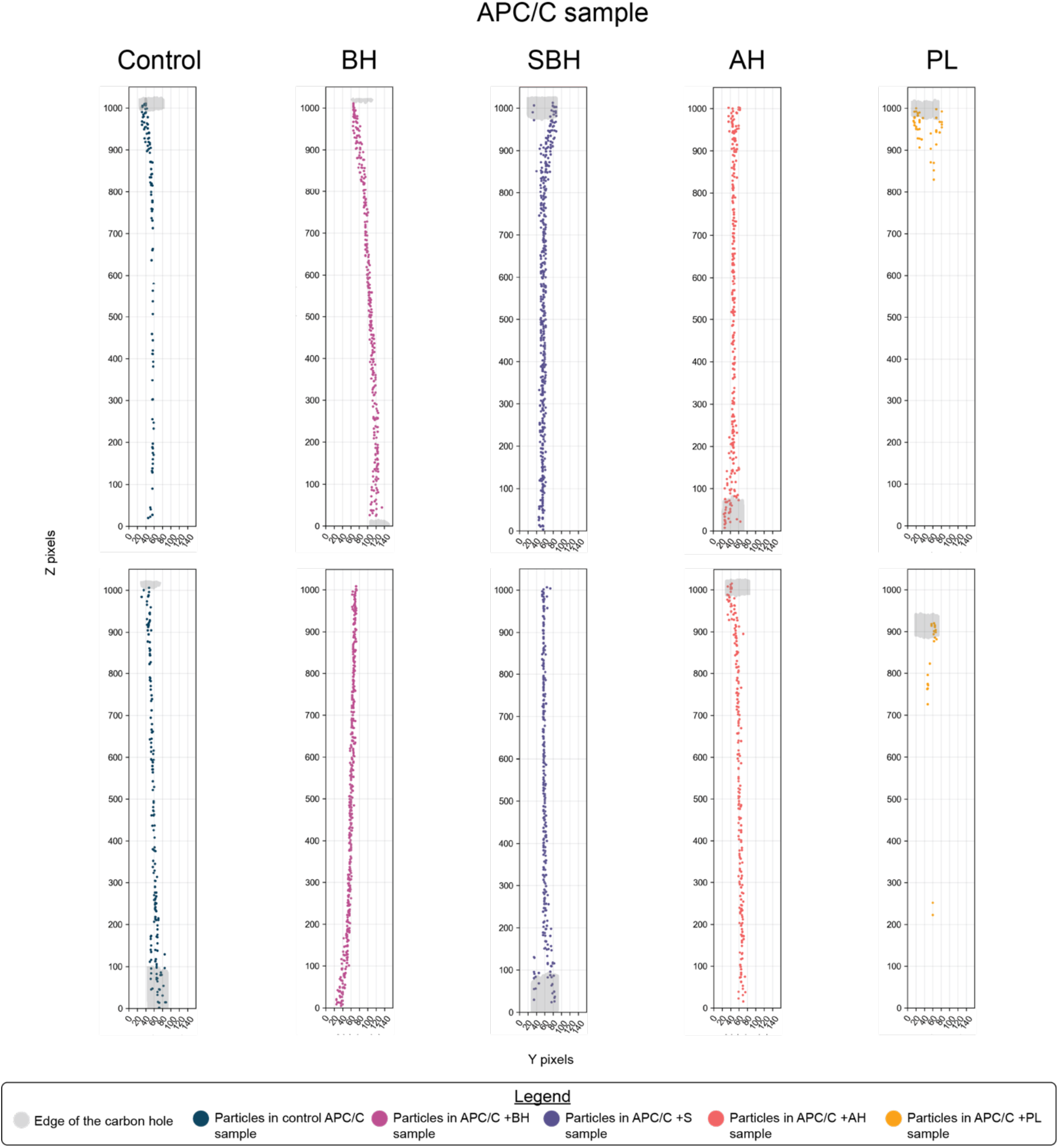
Picked tomograms. More picked tomograms of APC/C samples. The control sample particles are shown in dark blue, BH in pink, SBH in purple, AH in orange and PL in yellow. The carbon edge is shown in a transparent grey.

**Supplementary Figure 11:**
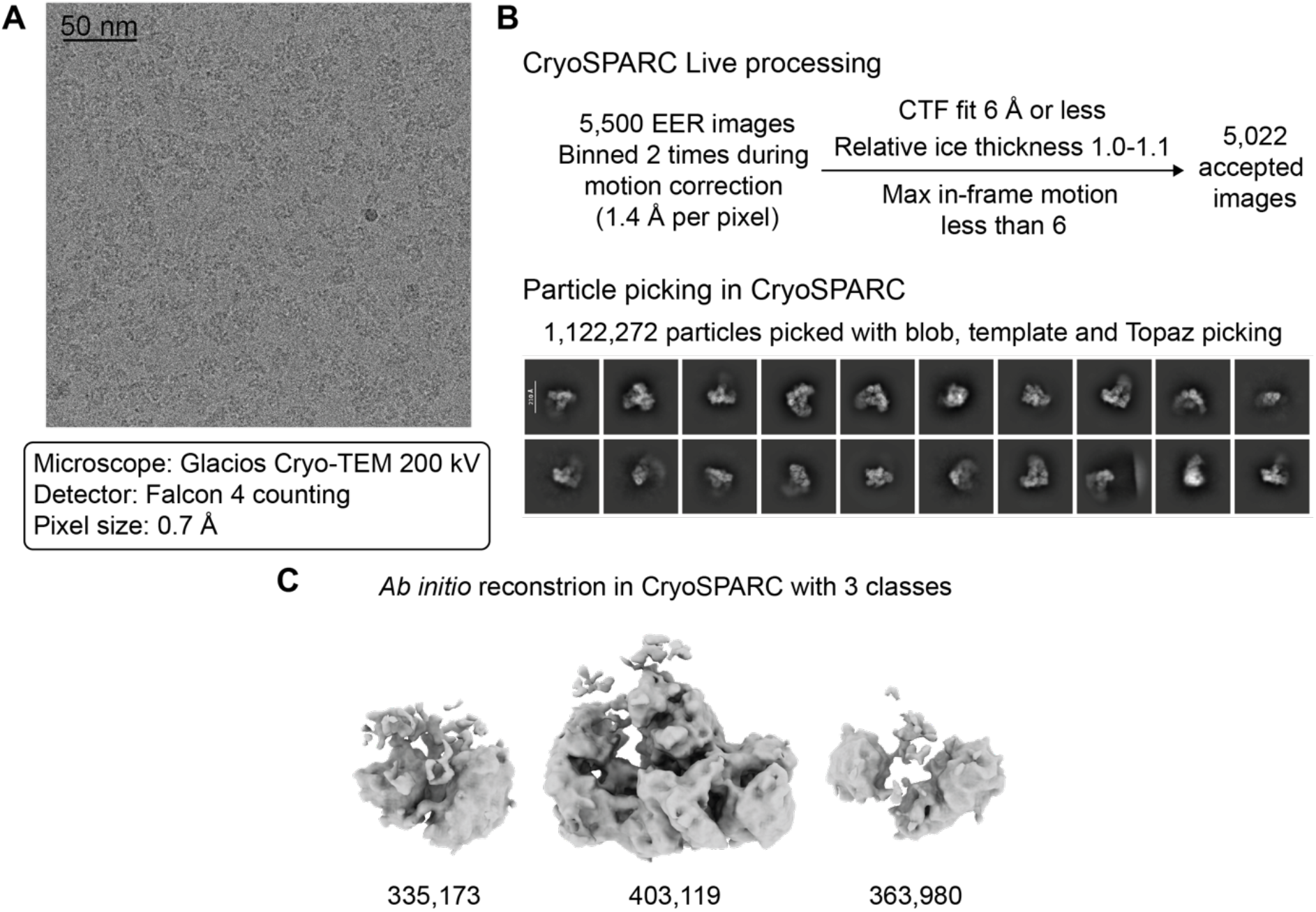
Cryo-EM analysis of the Control APC/C sample. (**A**) Representative micrograph and data collection parameters. (**B**) CryoSPARC processing for preprocessing micrographs and 2D particle picking. (**C**) Workflow of producing 3D reconstruction from *ab initio* in CryoSPARC.

**Supplementary Figure 12:**
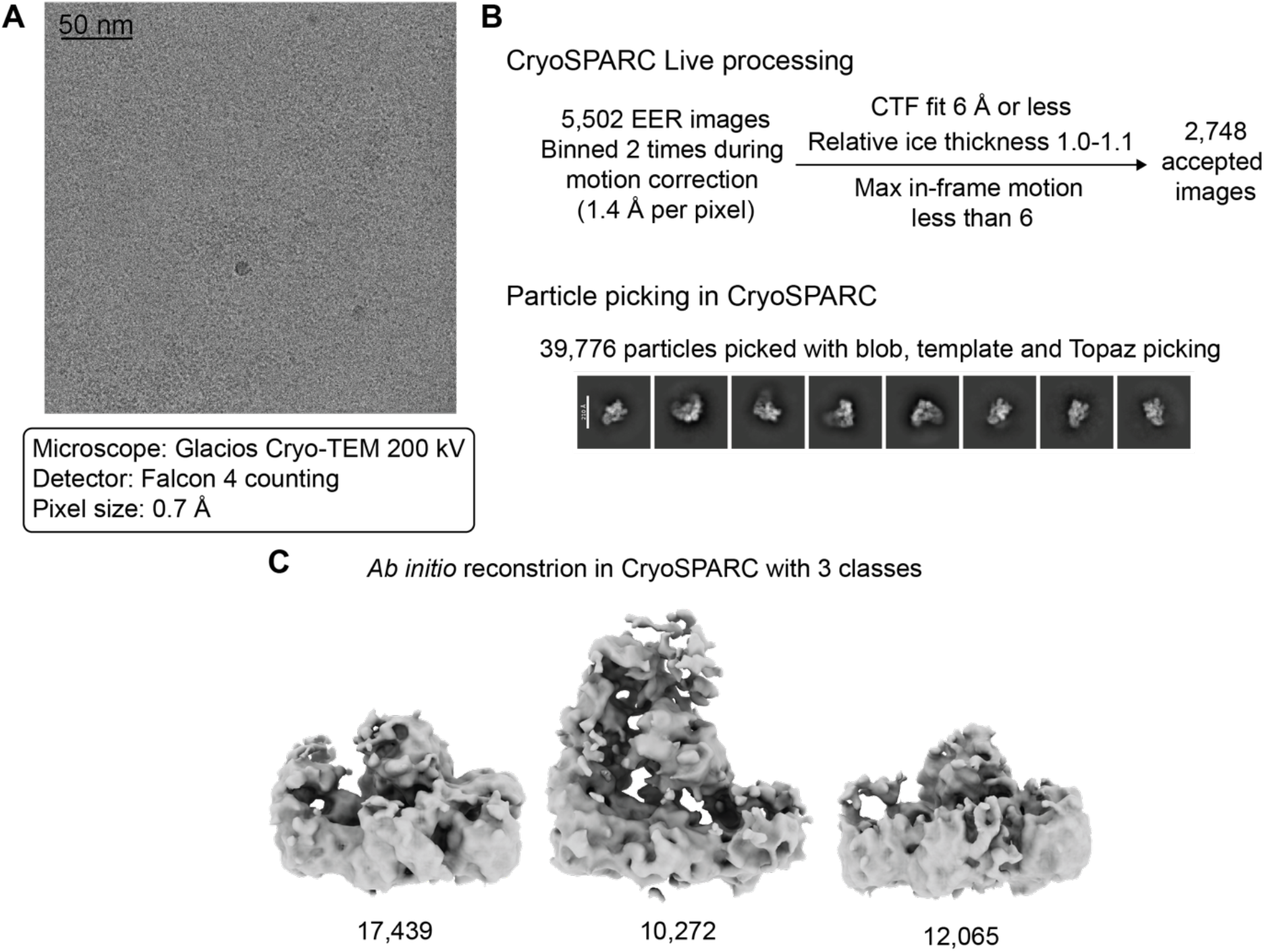
Cryo-EM analysis of the APC/C sample containing PL cryo-tide. (**A**) Representative micrograph and data collection parameters. (**B**) CryoSPARC processing for preprocessing micrographs and 2D particle picking. (**C**) *Ab initio* produced in CryoSPARC, with particle numbers listed.

**Supplementary Figure 13:**
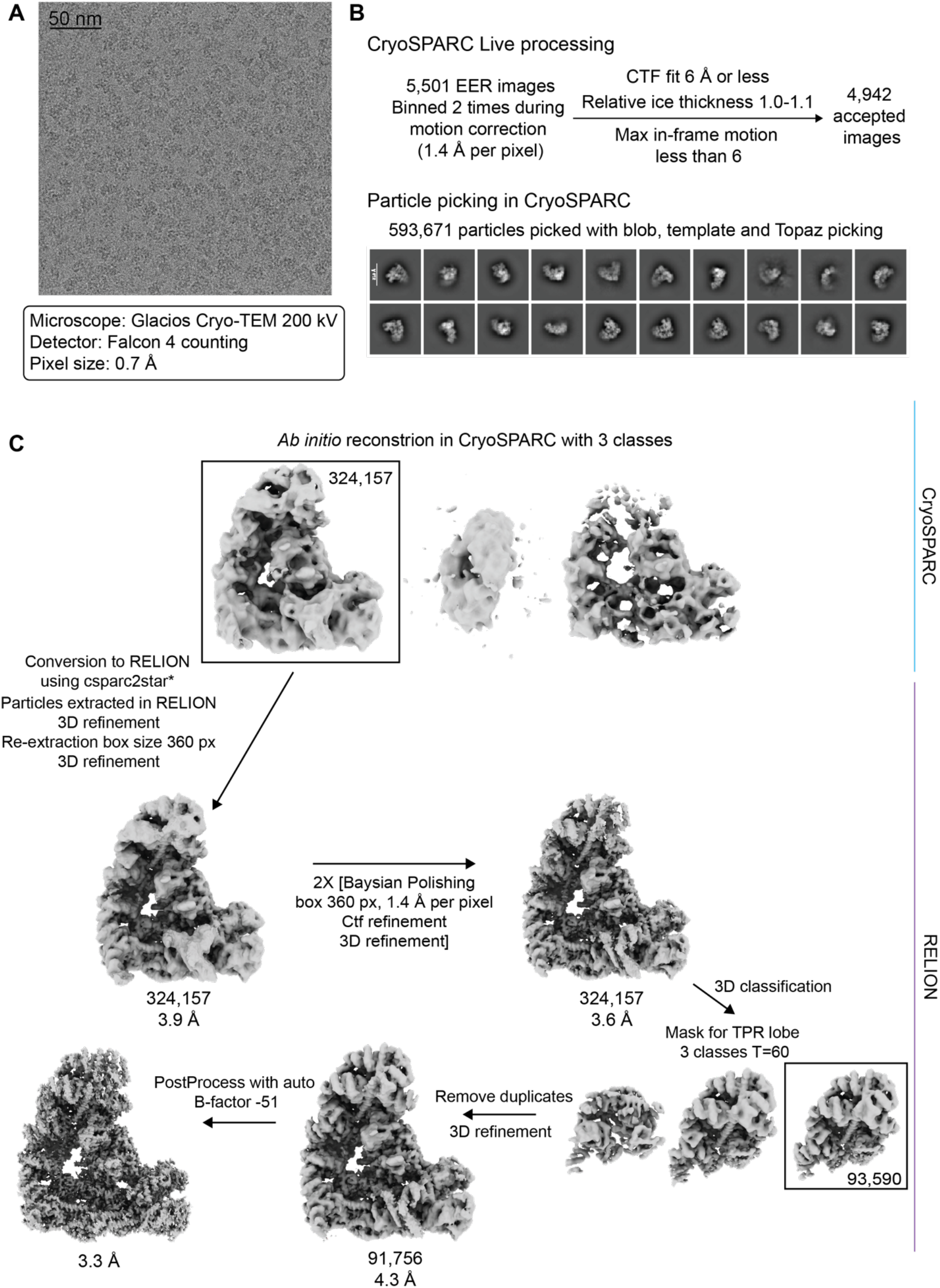
Cryo-EM analysis of the APC/C sample containing BH cryo-tide. (**A**) Representative micrograph and data collection parameters. (**B**) CryoSPARC processing for preprocessing micrographs and 2D particle picking. (**C**) Workflow of the key jobs for building the final 3D reconstructing. From *ab initio* produced in CryoSPARC, into RELION. Particles numbers displayed under maps. The conversion from CryoSPARC to RELION was done using csparc2star ^81^.

**Supplementary Figure 14:**
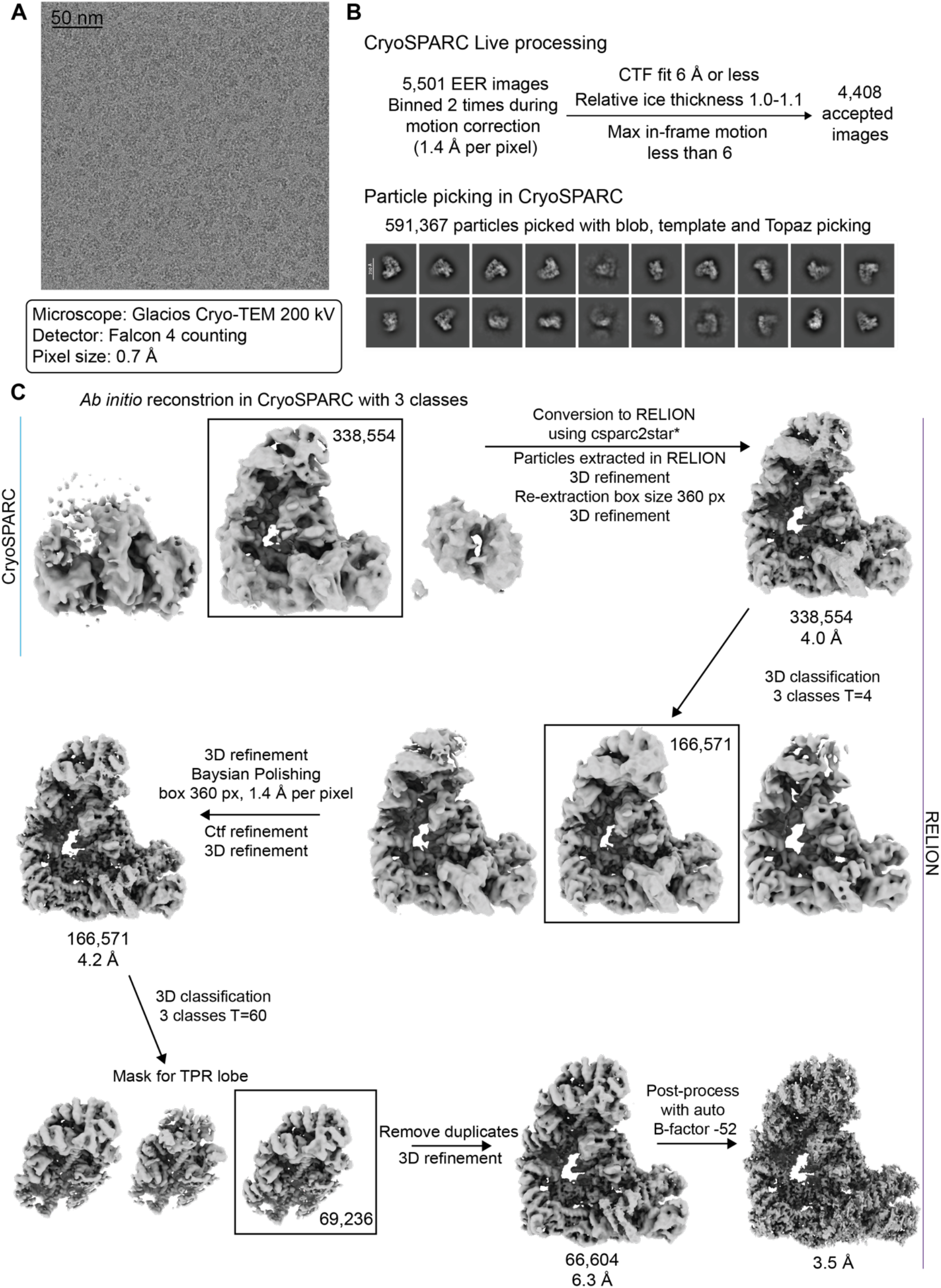
Cryo-EM analysis of the APC/C sample containing SBH cryo-tide. (**A**) Representative micrograph and data collection parameters. (**B**) CryoSPARC processing for preprocessing micrographs and 2D particle picking. (**C**) Workflow of the key jobs for building the final 3D reconstructing. From *ab initio* produced in CryoSPARC, into RELION. Particles numbers displayed under maps. The conversion from CryoSPARC to RELION was done using csparc2star ^81^.

**Supplementary Figure 15:**
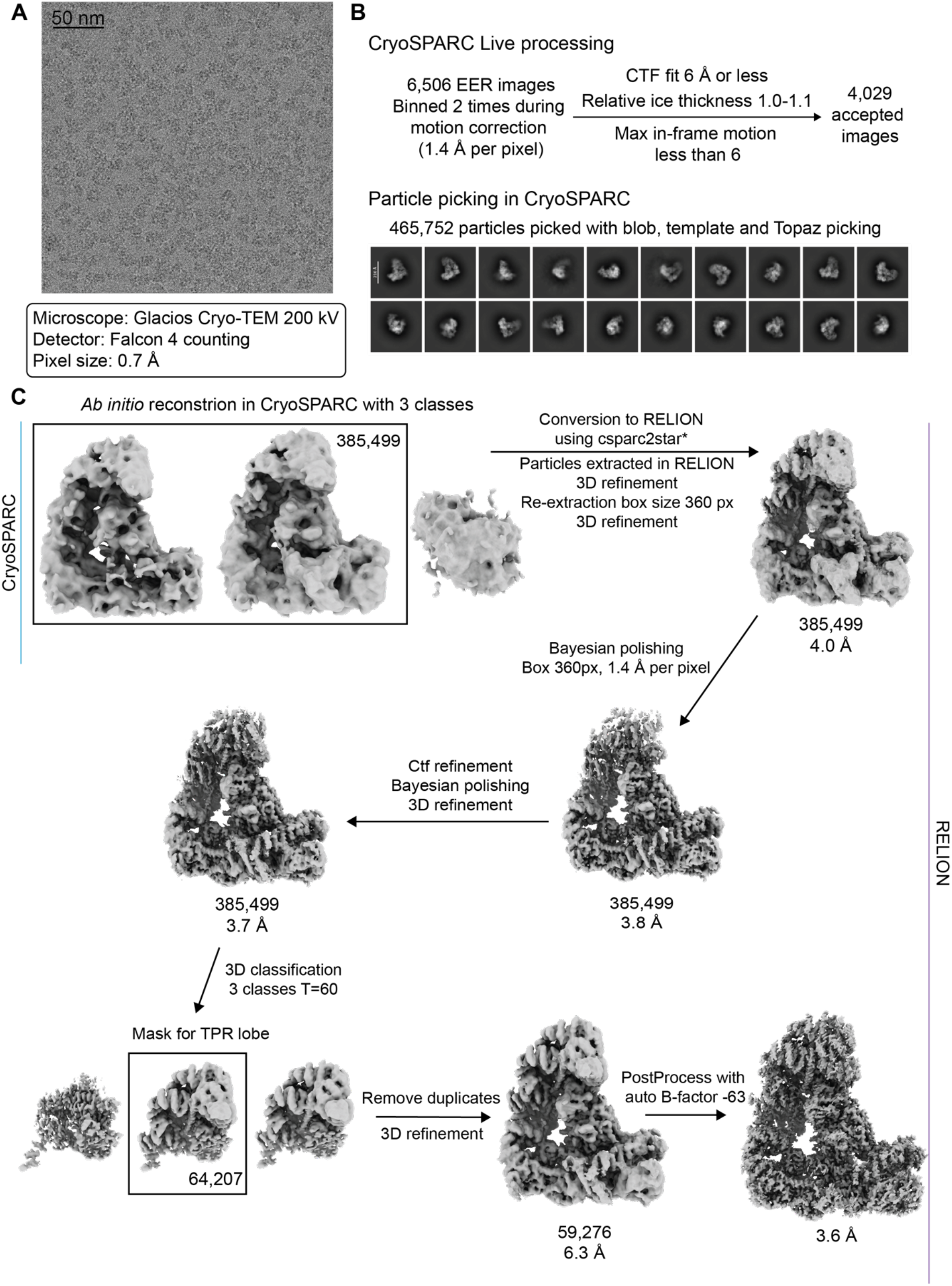
Cryo-EM analysis of the APC/C sample containing AH cryo-tide. (**A**) Representative micrograph and data collection parameters. (**B**) CryoSPARC processing for preprocessing micrographs and 2D particle picking. (**C**) Workflow of the key jobs for building the final 3D reconstructing. From *ab initio* produced in CryoSPARC, into RELION. Particles numbers displayed under maps. The conversion from CryoSPARC to RELION was done using csparc2star ^81^.

**Supplementary Figure 16:**
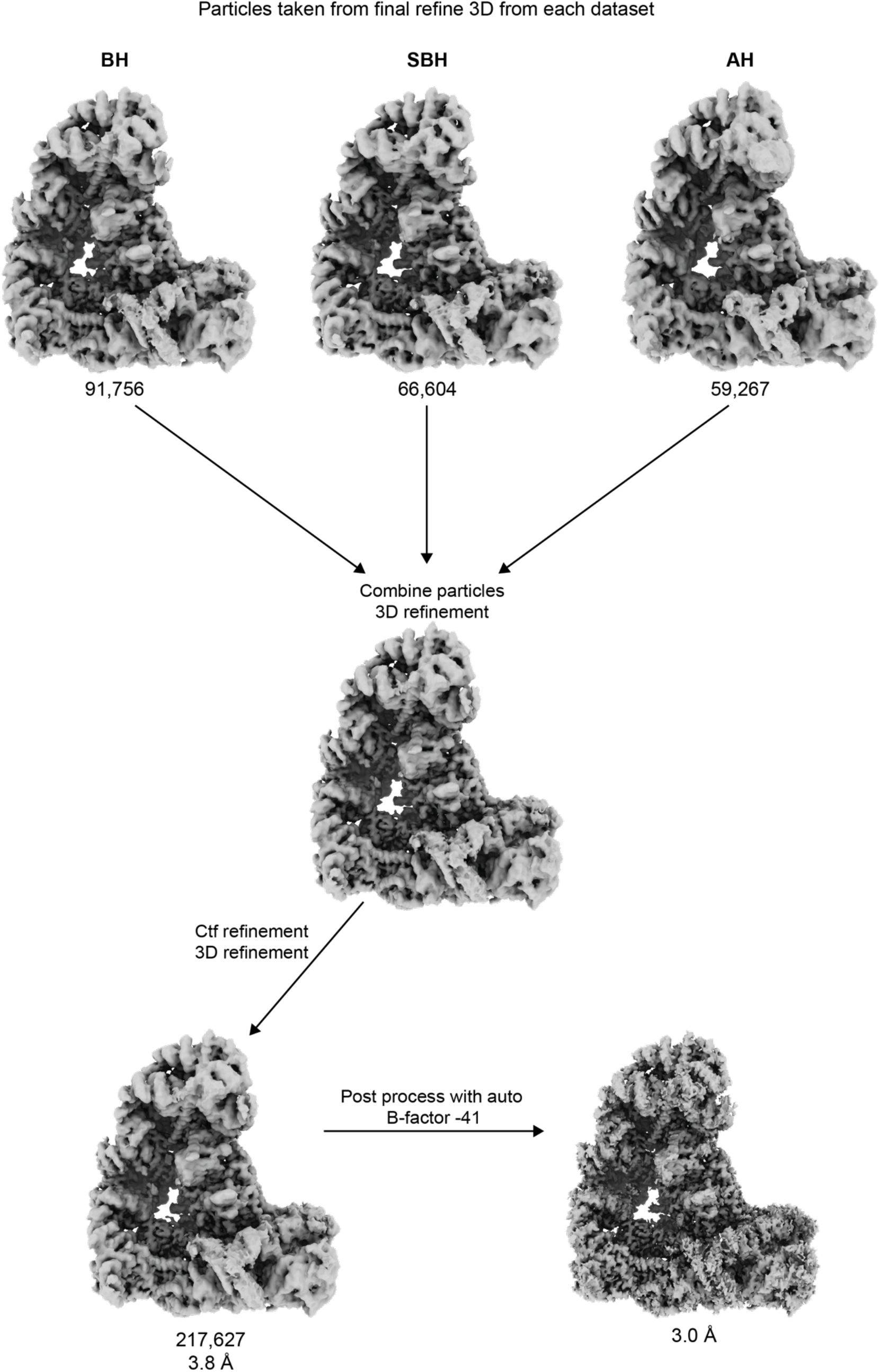
Image processing for combined cryo-tide dataset. The workflow from combining the final 3D refinements for each of the individual cryo-tide datasets through to the final refinement and postprocessed map of the combined dataset.

**Supplementary Figure 8:**
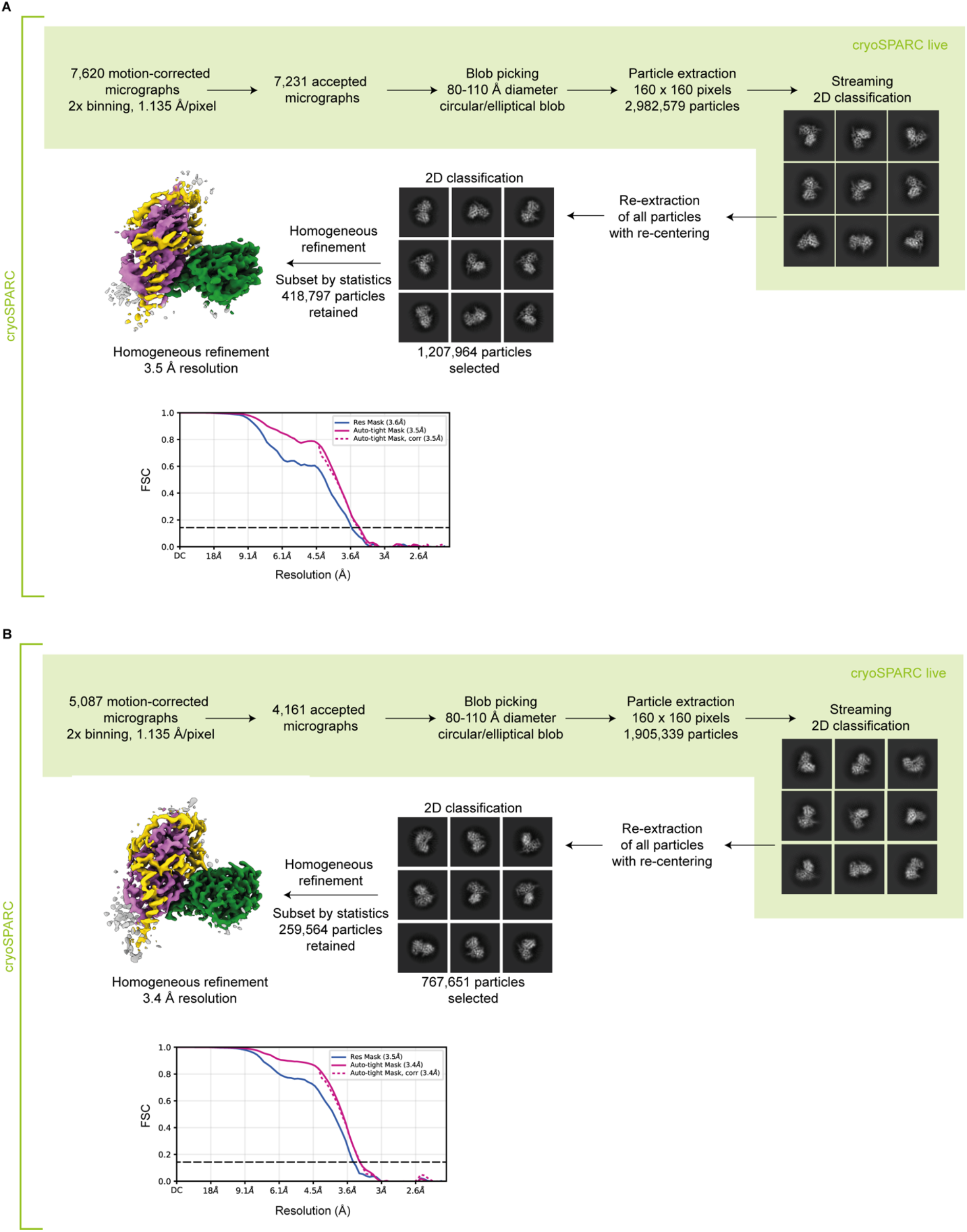
Image processing for cryo-EM samples of the CDK11:cyclin L:SAP30BP complex. (**A**) Image processing workflow for dataset without cryo-tide. (**B**) Image processing workflow for dataset with cryo-tide.

**Supplementary Table 1:** Cryo-EM data collection statistics for APC/C specimens.

| **Data collection and processing** | **Combined** | **Control** | **BH** | **SBH** | **AH** | **PL** |
| --- | --- | --- | --- | --- | --- | --- |
| Nominal magnification | N/A | 165,000 | 165,000 | 165,000 | 165,000 | 165,000 |
| Calibrated magnification | N/A | 200,000 | 200,000 | 200,000 | 200,000 | 200,000 |
| Acceleration voltage (kV) | N/A | 200 | 200 | 200 | 200 | 200 |
| Electron exposure (e<sup>-</sup>/Å<sup>2</sup>) | N/A | 60 | 60 | 60 | 60 | 60 |
| Defocus range (μM) | N/A | -0.6 to -1.6 | -0.6 to -1.6 | -0.6 to -1.6 | -0.6 to -1.6 | -0.6 to -1.6 |
| Pixel size (Å) | N/A | 0.7 | 0.7 | 0.7 | 0.7 | 0.7 |
| Symmetry imposed | N/A | C1 | C1 | C1 | C1 | C1 |
| Initial particle images (no.) | N/A | 1,122,272 | 593,671 | 591,367 | 465,752 | 39,776 |
| Final particle images (no.) | 217,627 | N/A | 91,756 | 66,604 | 59,276 | N/A |
| Map resolution (Å) | 3.0 | N/A | 3.3 | 3.5 | 3.6 | N/A |
| FSC threshold | 0.143 | N/A | 0.143 | 0.143 | 0.143 | N/A |

**Supplementary Table 2:** Cryo-EM data collection statistics for CDK11:cyclin L:SAP30BP specimens.

| **Data collection and processing** | **Control** | **BH** |
| --- | --- | --- |
|  | **EMD-59992** | **EMD-59987** |
| Nominal magnification | 240,000 | 240,000 |
| Calibrated magnification | 246,696 | 246,696 |
| Acceleration voltage (kV) | 200 | 200 |
| Electron exposure (e<sup>-</sup>/Å<sup>2</sup>) | 60 | 60 |
| Defocus range (μM) | -0.5 to -1.8 | -0.5 to -1.8 |
| Pixel size (Å) | 0.5675 | 0.5675 |
| Symmetry imposed | C1 | C1 |
| Initial particle images (no.) | 2,982,579 | 1,905,339 |
| Final particle images (no.) | 418,797 | 259,564 |
| Map resolution (Å) | 3.5 | 3.4 |
| FSC threshold | 0.143 | 0.143 |

**Supplementary Table 3:** Cryo-EM refinement and validation statistics.

| **Refinement** | **Combined** |
| --- | --- |
| Initial model used (PDB code) | 8PKP |
| Model resolution (Å) | 3.2 |
| FSC threshold | 0.5 |
| Map sharpening B factor (Å<sup>2</sup>) | -41 |
| <u>Model composition</u> |  |
| Nonhydrogen atoms | 62538 |
| Protein residues | 7935 |
| Nucleotides | 0 |
| Ligands | ZN:4 |
| <u>B factors (Å<sup>2</sup>)</u> |  |
| Protein (min/max/mean) | 48.76/376.00/129.85 |
| Ligand | 376.26/413.44/391.33 |
| <u>R.M.S deviations</u> |  |
| Bond lengths (Å) | 0.002 |
| Bond angles (°) | 0.548 |
| **Validation** |  |
| MolProbity score | 1.86 |
| Clashscore | 6.35 |
| Poor rotamers (%) | 2.19 |
| <u>Ramachandran plot</u> |  |
| Favour (%) | 96.30 |
| Allowed (%) | 3.60 |
| Disallowed (%) | 0.10 |

